# Absolute measurement of fast and slow neuronal signals with fluorescence lifetime photometry at high temporal resolution

**DOI:** 10.1101/2025.01.10.632162

**Authors:** Bart Lodder, Tarun Kamath, Ecaterina Savenco, Berend Röring, Michelle Siegel, Julie Chouinard, Suk Joon Lee, Caroline Zagoren, Paul Rosen, Roger Adan, Lin Tian, Bernardo L. Sabatini

## Abstract

The concentrations of extracellular and intracellular signaling molecules, such as dopamine and cAMP, change over both fast and slow timescales and impact downstream pathways in a cell-type specific manner. Fluorescence sensors currently used to monitor such signals *in vivo* are typically optimized to detect fast, relative changes in concentration of the target molecule. They are less well suited to detect slowly-changing signals and rarely provide absolute measurements of either fast and slow signaling components. Here, we developed a system for fluorescence lifetime photometry at high temporal resolution (FLIPR) that utilizes frequency-domain analog processing to measure the absolute fluorescence lifetime of genetically-encoded sensors at high speed but with long-term stability and picosecond precision in freely moving mice. We applied FLIPR to investigate dopamine signaling in two functionally distinct regions in the striatum, the nucleus accumbens core (NAC) and the tail of striatum (TOS). We observed higher tonic dopamine levels at baseline in the TOS compared to the NAC and detected differential and dynamic responses in phasic and tonic dopamine to appetitive and aversive stimuli. Thus, FLIPR enables simple monitoring of fast and slow time-scale neuronal signaling in absolute units, revealing previously unappreciated spatial and temporal variation even in well-studied signaling systems.

## Introduction

The development of genetically-encoded fluorescent indicators (GEFI) has greatly enhanced our capabilities to investigate neuronal signaling. Many neurotransmitters, neuromodulators, and intracellular signals can be measured using GEFIs ^1–7^. Moreover, GEFIs can be genetically targeted to specific neuronal populations and, when combined with fiber photometric approaches, monitored in freely moving animals^8–10^. Most GEFIs are designed to change brightness upon substrate binding and thus are typically used with systems that measure fluorescence intensity over time.

Ideally GEFIs would report neuronal signals in absolute, physiologically relevant units – such as numbers of action potentials or the concentration of a neuromodulator or second messenger. This is typically impractical because fluorescence intensity is a property of both the state of individual sensors (typically bound or unbound to the molecule they detect) and bulk properties such as sensor concentration, which vary across cells, sites and animals^11,12^. For these reasons, fluorescence intensity is usually converted into relative units by dividing by baseline intensity (dF/F), signal variance (z-scoring), or a ligand-insensitive signal such as the isosbestic point ^8,10,13^. These normalized signals can still be dynamically impacted by hemodynamic artifacts caused by the differential absorption of excitation and emission light by blood, by animal movement, and by differential photobleaching of signal (GEFI) and background (e.g., autofluorescence) ^11,12,14–17^. Such substrate-independent changes in fluorescence intensity also limit the timescale over which quantitative comparisons can be made – fluctuations in fluorescence intensity due to slow changes in substrate concentration are difficult to distinguish from bleaching or hemodynamic and motion artifacts ^10^. Furthermore, photobleaching typically requires relative measures to be calculated over a rolling window, which limits the detection of signals that occur more slowly than a small fraction of the window length ^10,13^. As a consequence, standard fluorescence intensity measurements can capture rapid transient changes of for instance dopamine but cannot capture changes in slow tonic dopamine.

An alternative measurement modality is to monitor state-dependent changes in the fluorescence lifetime, instead of the intensity, of specially designed GEFIs. Fluorescence lifetime is the average time after excitation that a fluorophore takes to release a photon and is an intrinsic property of each GEFI molecule ^12^. As such, it is not dependent on sensor concentration or changes in intensity (as long as the sensor fluorescence is substantially brighter than autofluorescence) ^9,18,19^ . Therefore, fluorescence lifetime measurements are insensitive to many of the caveats that limit absolute intensity imaging, including variability in expression, photobleaching, and hemodynamic and motion artifacts ^9,18,19^. Altogether, this means that fluorescence lifetime can be expressed in absolute units of time (typically picoseconds (ps) or nanoseconds (ns)) and used to monitor signals across a wide range of time scales without post-processing or artifact-imposed limitations on the ability to measure slow signals.

The widespread use of fluorescence lifetime to monitor neuronal signals *in vivo* has been hampered by three main factors. First, fluorescence lifetime is typically measured with time correlated single photon counting (TCSPC) systems that record the lifetime of one GEFI molecule at a time. Measurements are combined to form a histogram that can be fitted with single or multi-exponential curves to calculate the fluorescence lifetime of a single or multiple fluorescence lifetime agents, respectively ^17,20,21^. Collecting sufficient individual emission events to achieve acceptable signal-to-noise ratios is, due to hardware constraints, typically slow, limiting the temporal resolution to ∼1 Hz ^17,20,21^. Second, existing fluorescence lifetime systems are complex and expensive, often relying on highly specialized and fragile equipment ^9,10^. Third, most GEFIs do not exhibit state-dependent changes in lifetime, such that only a few fluorescence lifetime sensors are currently available to measure neuronal signals ^3,17,22–27^.

With the goal of facilitating the high-speed absolute measurement of neuronal signals *in vivo* in absolute units and over a wide range of timescales, we designed a high-speed, precise, low-cost, robust, and easy to use fluorescence lifetime photometry (FLIP) system, termed fluorescence lifetime photometry at high temporal resolution (FLIPR). FLIPR relies on frequency domain analog processing to calculate fluorescence lifetime at picosecond precision *in vivo*. We show that FLIPR enables accurate fluorescence lifetime detection in freely moving mice, preserving signals over a wide dynamic range from baseline (DC) to rapid (10-1000 Hz) fluctuations. Furthermore, improvements in the FLIPR analog processing electronics allow for artifact-free measurement across a large fluorescence intensity range. We demonstrate the utility of FLIPR by validating and applying a lifetime dopamine sensor to concurrently measure phasic and tonic dopamine in behaving mice. This reveals absolute differences in tonic dopamine across brain regions and changes in absolute phasic and tonic dopamine during appetitive and aversive stimuli.

## Results

### Design of fluorescence lifetime photometry at high temporal resolution (FLIPR)

To enable high speed and affordable fluorescence lifetime recordings in the brain of behaving animals, we developed FLIPR, which builds on previous work in fluorescence lifetime photometry and microscopy ^8,9,20,28^ (**Figure 1** and **S1**). The excitation and emission paths are similar to those used in typical fiber photometry system. A 50 MHz pulsed laser provides excitation light, which is intensity modulated using an acousto-optic modulator and focused into a multimode optical fiber with appropriate elements. The optical fiber couples end-to-end with an optical fiber stub implanted in the brain of a mouse. Emitted fluorescence returns through the same path and, on exiting from the optical fiber, is collimated, separated from the excitation light with a dichroic and interference filters, and directed to a photomultiplier tube (PMT).

**Figure 1.**
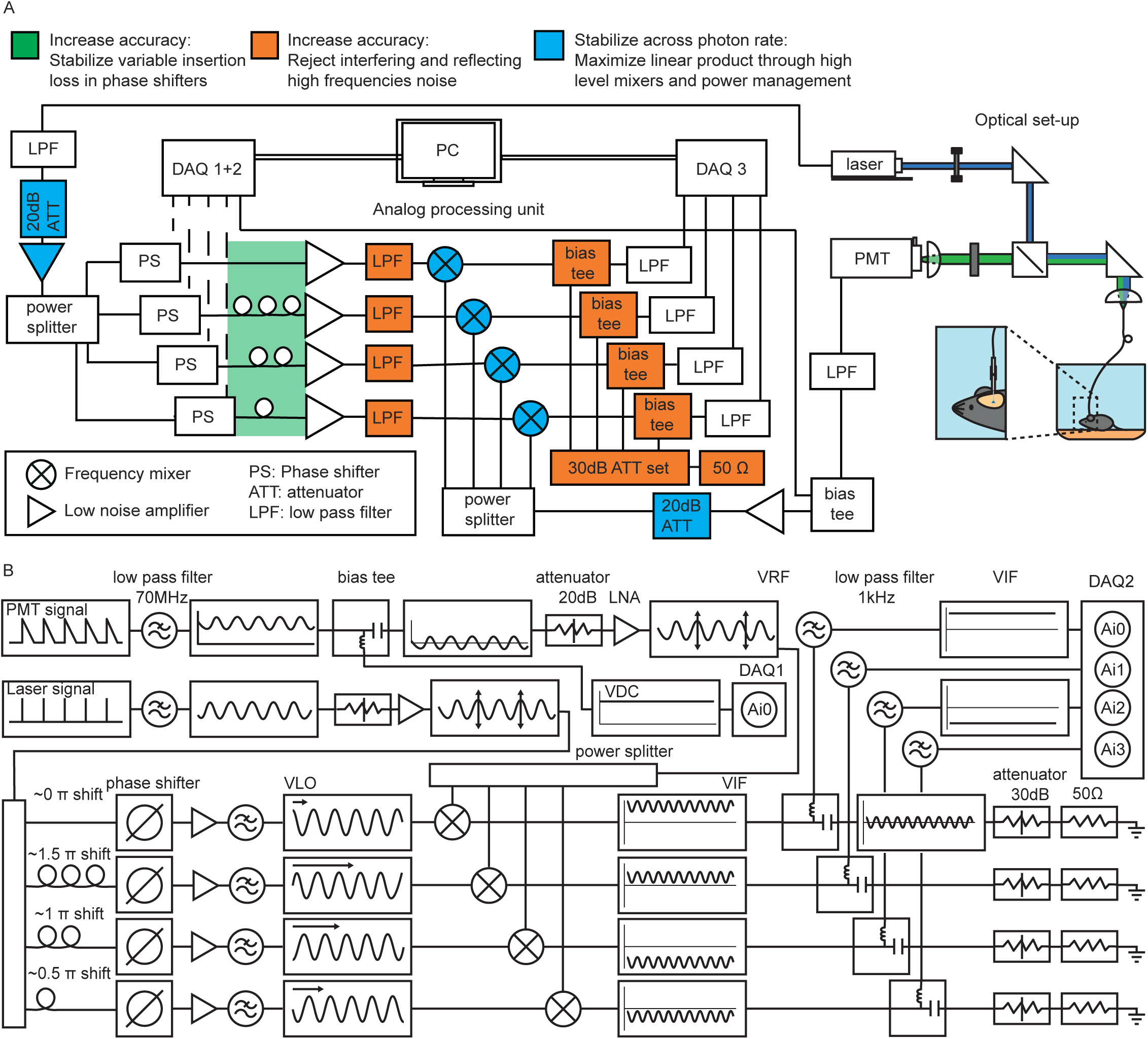
FLIPR system diagram **A,** FLIPR consists of an optical path (right) and analog processing unit (left). Excitation light is provided by a 50 MHz pulsed 473 nm laser whose output is focused into a fiber optic that couples to a fiber optic stub implanted in the brain. The same fiber optic collects green fluorescence which is separated from the excitation light using a dichroic filter and directed to a PMT. The analog processing unit performs high-speed phase detection using phase shifting and mixing circuitry. Blue, green and orange highlight critical improvements to stabilize and increase the accuracy of fluorescence lifetime calculations across power levels. LPF: low pass filter; ATT: electrical attenuator; LNA: low noise amplifier; PS: phase shifter; DAQ: data acquisition system; TA: transimpedance amplifier; PMT: photon multiplier tube. **B**, Schematic of signal processing in the FLIPR computing unit. In the top left, a schematized signal from the PMT and laser reference signal is passed through a low pass filter to extract the 50 MHz modulated signal. The PMT signal is split by a bias tee into a high frequency signal (50 MHz) and DC or intensity (VDC) signal. The high frequency PMT signal (VRF) is power adjusted by an attenuator and low noise amplifier (LNA), and passed into the frequency mixers. The laser reference signal is power adjusted by an attenuator and LNA and split into 4 different channels by a power splitter. The 4 channels are shifted by 0, 0.5, 1 and 1.5 π through physical delay lines and controllable phase shifters, amplified using LNAs and low pass filtered at 70 MHz. The laser reference signal (VLO) channels are passed into the frequency mixers, where multiplication with VRF produces a low frequency (DC-kHz) and high frequency (∼100 MHz) VIF signal. High frequency signals are filtered out and absorbed through bias tees, attenuators and 50 Ohm resistors. Low frequency information is low pass filtered at 1 kHz. All relevant signals are collected by a data acquisition system (DAQ). VLO: local oscillator voltage; VRF: radio frequency voltage; VIF: intermediate frequency voltage. VDC: direct current voltage.

FLIPR relies on an analog frequency domain fluorescence lifetime approach, wherein the fluorescence lifetime is calculated using an analog phase and intensity detection system (**Figure 1A,B** and **S1**). The fluorescence lifetime is calculated by comparing the phase difference of the PMT signal and the reference laser pulse. The electrical reference laser pulse signal is cleaned up with a low pass filter (70 MHz corner frequency) and referred to as the local oscillator voltage or VLO. The VLO is split into 4 channels that are shifted by 0, 0.5, 1 and 1.5 π radians with the zero phase shift being calibrated at 0 ns (or instantaneous lifetime) using a fluorescent lifetime standard. The PMT output is converted to voltage using an internal transimpedance amplifier, low-pass filtered at 70 MHz, and separated into a DC component (VDC, which is the fluorescence intensity signal) and an AC signal (radio frequency voltage, VRF). The VRF is split into 4 copies, each of which is mixed with one of the phase-shifted reference signals. The mixers multiply each phase shifted VLO, corresponding to the laser reference, and the VRF, corresponding to the PMT output, producing outputs that depend on the relative phases and amplitudes of the VRF and VLO signals. The four signals (intermediate frequency voltage, VIF) are passed through 1 kHz low pass filters and, along with the intensity VDC signal, captured using a data acquisition system. The VIF and VDC voltage allow for real-time calculation of lifetime and phasor components (see methods). Phasor analysis can be used to determine if the fluorescence follows a single exponential or multi-exponential decay: lifetimes with a single exponential decay have phasor components located on a semi-circle termed the universal circle (centered at g = 0.5, s = 0, radius = 0.5), whereas lifetimes with a multi-exponential decay fall outside of the universal circle ^29^. These signals are sampled by the data acquisition system at the temporal resolution set by the dynamics of neuronal signaling (typically <1 kHz) as opposed to by the fluorescence lifetime (∼10 GHz).

Although the principle of operation of the FLIPR system is straight forward and described previously ^20^, we made several crucial changes to the FLIPR analog frequency domain computing unit that improve the accuracy and power stability as necessary for use in a real laboratory setting.

First, the amplifiers and frequency mixers produce non-linear responses when the VRF approaches the 1dB compression point ^30^. This in turn leads to a channel specific reduction in voltage, which can cause substantial lifetime and phasor readout artifacts as a function of fluorescence intensity. To increase the stability of fluorescence lifetime readout across a broad intensity range, we used frequency mixers with a higher 1dB compression point. In addition, we added RF amplifiers and attenuators to the VLO and VRF paths. In the VLO path, we increased the VLO signal amplitude for optimal mixer performance whereas, in the VRF path, we reduced signal amplitude to achieve a more linear response and reduce the voltage fluctuations with fluorescence intensity changes. In order to accommodate the low VRF amplitudes, we selected a free standing DAQ unit with high voltage accuracy and stability.

Second, the shift of each of the four phase shifters needs to be set empirically using control voltages. However, the wide range of voltages necessary to produce 0 – 1.5 π radian shifts leads to differential insertion loss between the channels. The resulting different VLO magnitudes across channels imbalances the mixer products, introducing inaccuracies in the fluorescence lifetime estimation. To solve this problem, we accomplished the majority of the phase shift by altering the length of physical delay lines and used the phase shifters only to fine-tune the phase shift and calibrate the system. In this configuration, the difference between control voltage over the phase shifters is minimal and differences between VLO voltages are diminished.

Third, we observed several electrical reflections in the system that affected mixer function. The mixers output the sum and the differences of the VLO and RF frequencies, i.e. components at 0 and 100 MHz. Without the design features described below, the 100 MHz signal is reflected by a low pass filter and can re-enter the mixer where it is multiplied with the VLO ^30,31^. This creates new 50 and 150 MHz signals that are again reflected at the low pass filter. The interaction of these reflections with the VLO produces a low frequency signal (VIF_reflection_) that sums with the initial low frequency signal, introducing an artifact. Moreover, the amplitude of VIF_reflection_ dependents upon the VLO phase shifts and therefore is channel dependent. This also imbalances the mixer outputs and perturbs the lifetime readout. To solve this problem, we split the high frequency mixer outputs from the low frequency mixer outputs using bias tees and pass the high frequency signal through high dB absorbing attenuators and terminate them at 50 Ohm, eliminating the reflection. Finally, we also reject higher frequencies produced by the phase shifters by placing low pass filters before the mixers. Altogether, these changes greatly increase the accuracy and stability of the phase detection system.

To control the FLIPR system, acquire date and display fluorescence lifetime in real time, we developed a MATLAB based software, adapted from Sabatini lab electrophysiology acquisition software (**Figure S1**). The cost of building this new FLIPR system, excludes tools and a computer, is estimated ∼$38,000 (**Table 1**), with ∼50% of the cost being the pulse laser which can, in theory, be shared across multiple FLIPR systems. The total cost of building the previously published time-domain (TCSPC) FLIP system is estimated at ∼$68,000.

**Table 1.**
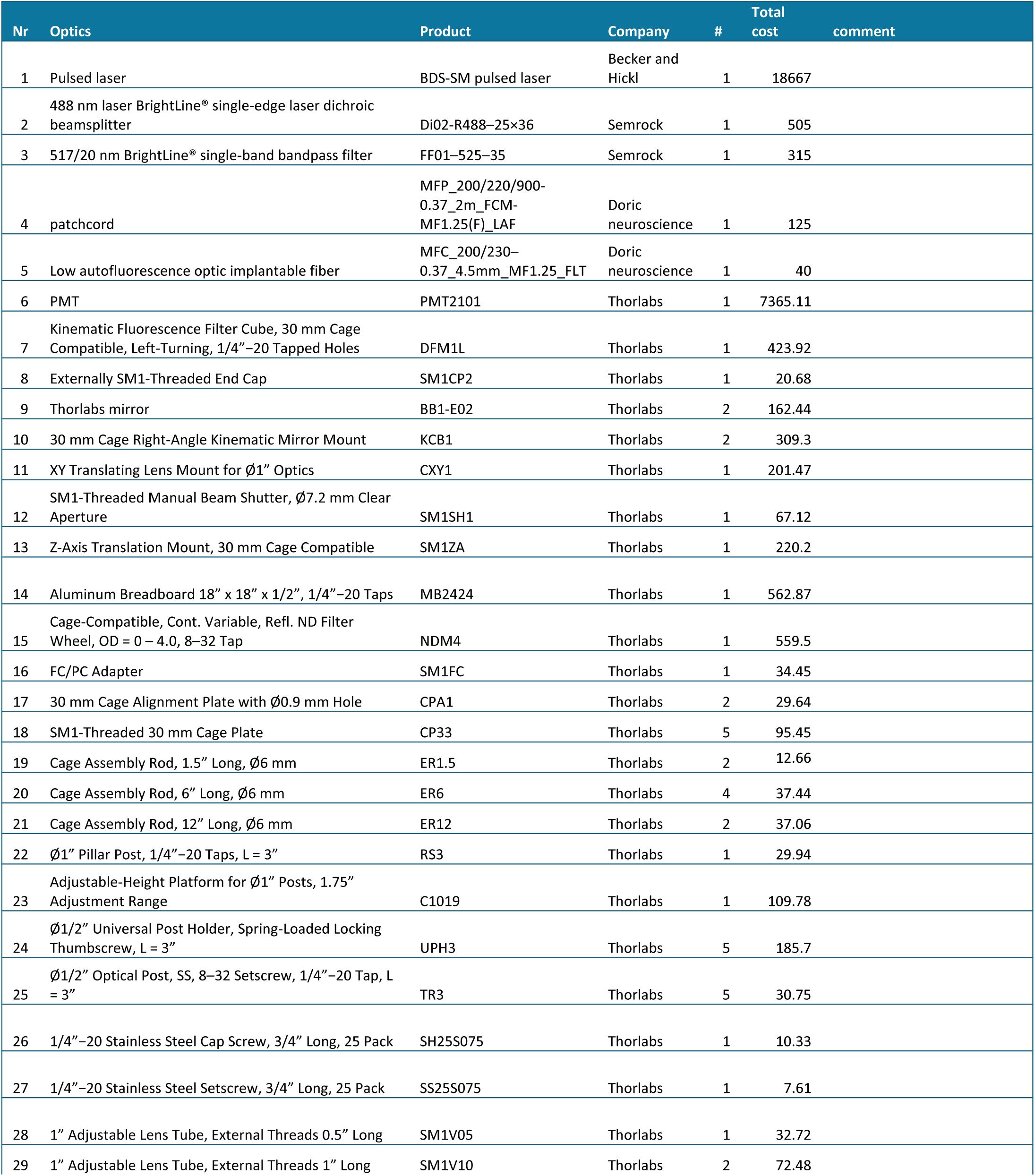
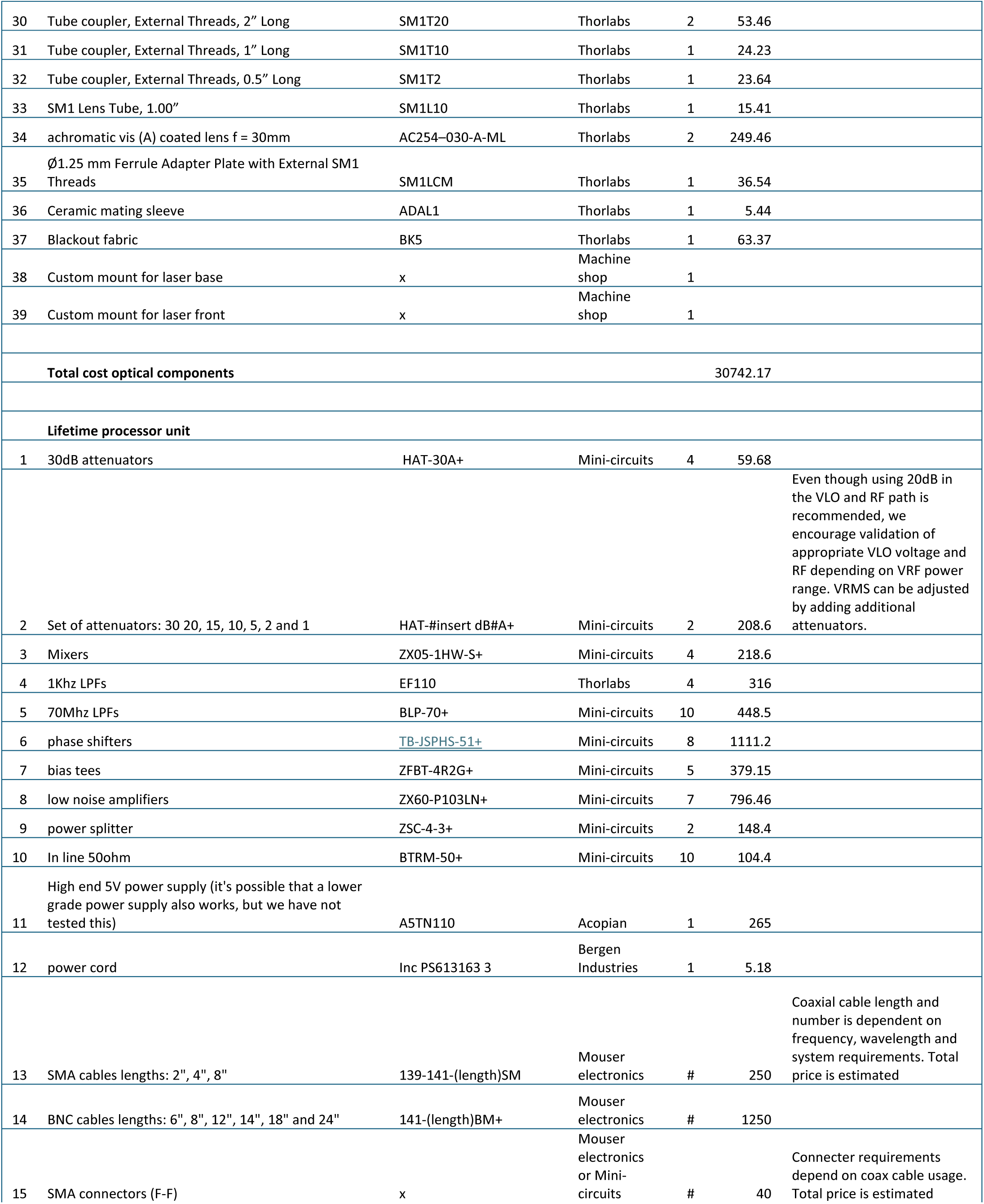
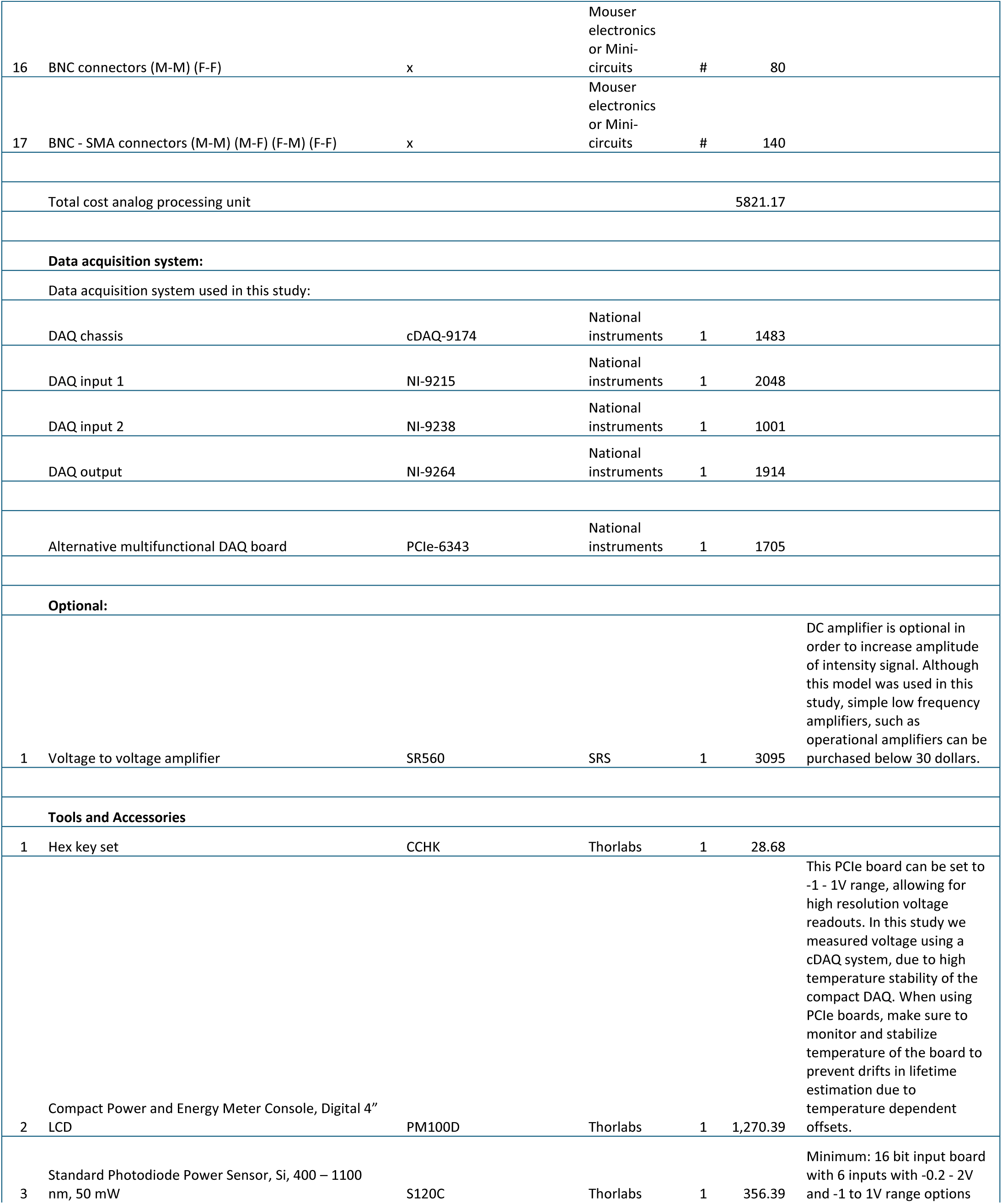
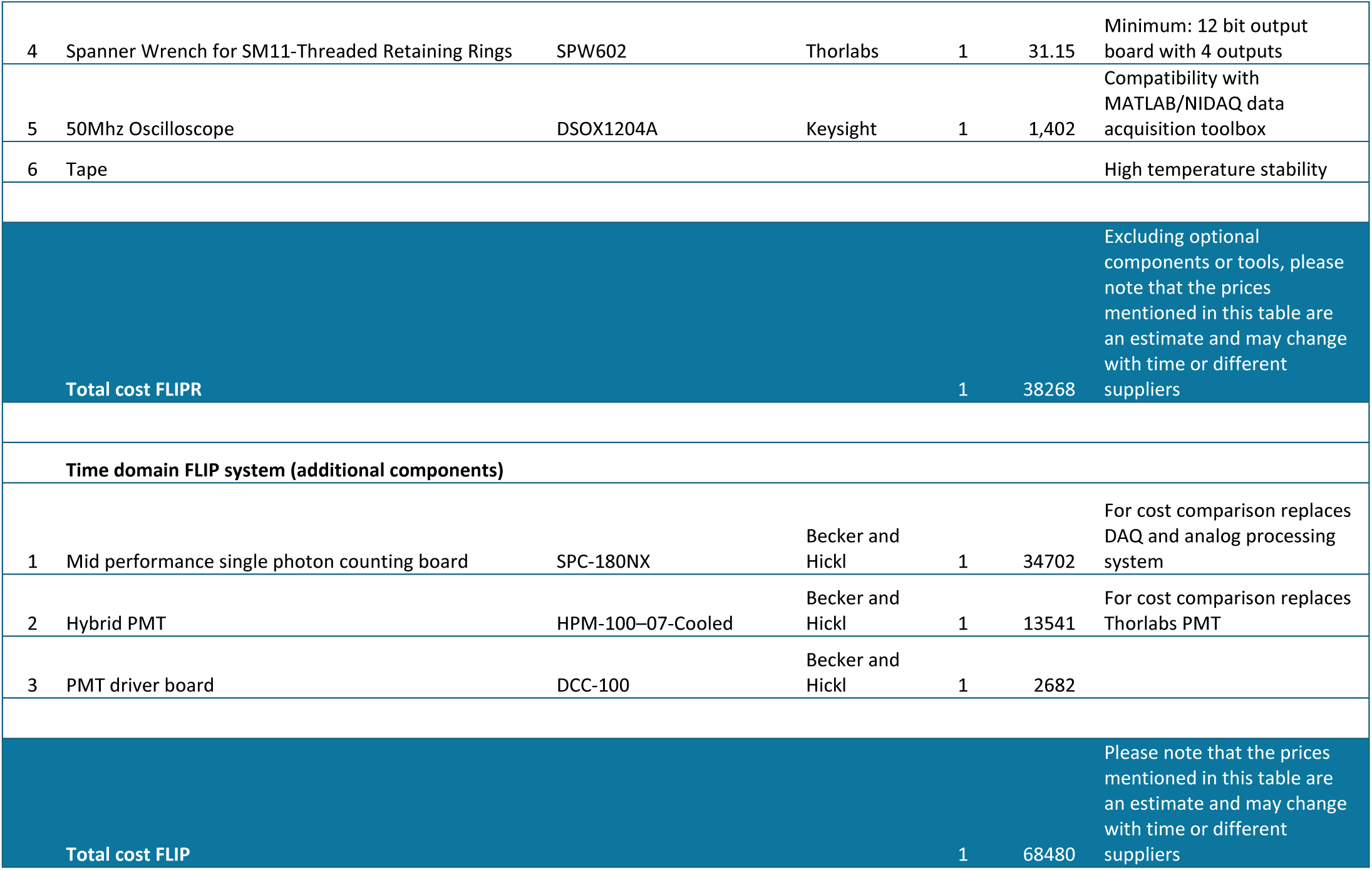

### *In vitro* validation

To validate the accuracy of the FLIPR system, we compared FLIPR to the time domain TCSPC based FLIP system that we previously built and used for biological discovery ^9,21,32^ (**Figure 2**). We built an optical layout that allowed switching between fluorescence lifetime measurements in either time (FLIP) or frequency (FLIPR) domain (**Figure S2**). We used this approach to compare the lifetimes of fluorescent compounds coumarin 6 (7.1 x 10^-8^ M in ethanol) , acridine orange (2.1 x 10^-7^ M in ethanol) and fluorescein (6.0 x 10^-6^ M in water) generated and collected with a fiber optic placed in a cuvette and measured with FLIP and FLIPR (**Figure 2A**). Estimated fluorescence lifetimes of coumarin 6, Aciridine orange and Fluorescein were essentially identical whether collected with FLIP or FLIPR (**Figure 2B**). In addition, to test the accuracy and linearity of FLIPR, the fluorescence lifetimes of mixtures of acridine orange and coumarin 6 at varying ratios were measured using FLIPR. As expected for the lifetime of a mixture of two fluorophores with different individual lifetimes, the weighted ratios of the acridine orange/coumarin 6 lifetimes and the measured fluorescence lifetime was linear (**Figure 2C**, R^2^ = 0.999). Furthermore, phasor components of coumarin 6, acridine orange and fluorescein fell on predicted positions on the universal half circle in the phasor plot (**Figure 2D**). Altogether, these data indicate that fluorescence lifetime measurements made with FLIPR are accurate and similar to those of conventional time-domain FLiP.

**Figure 2.**
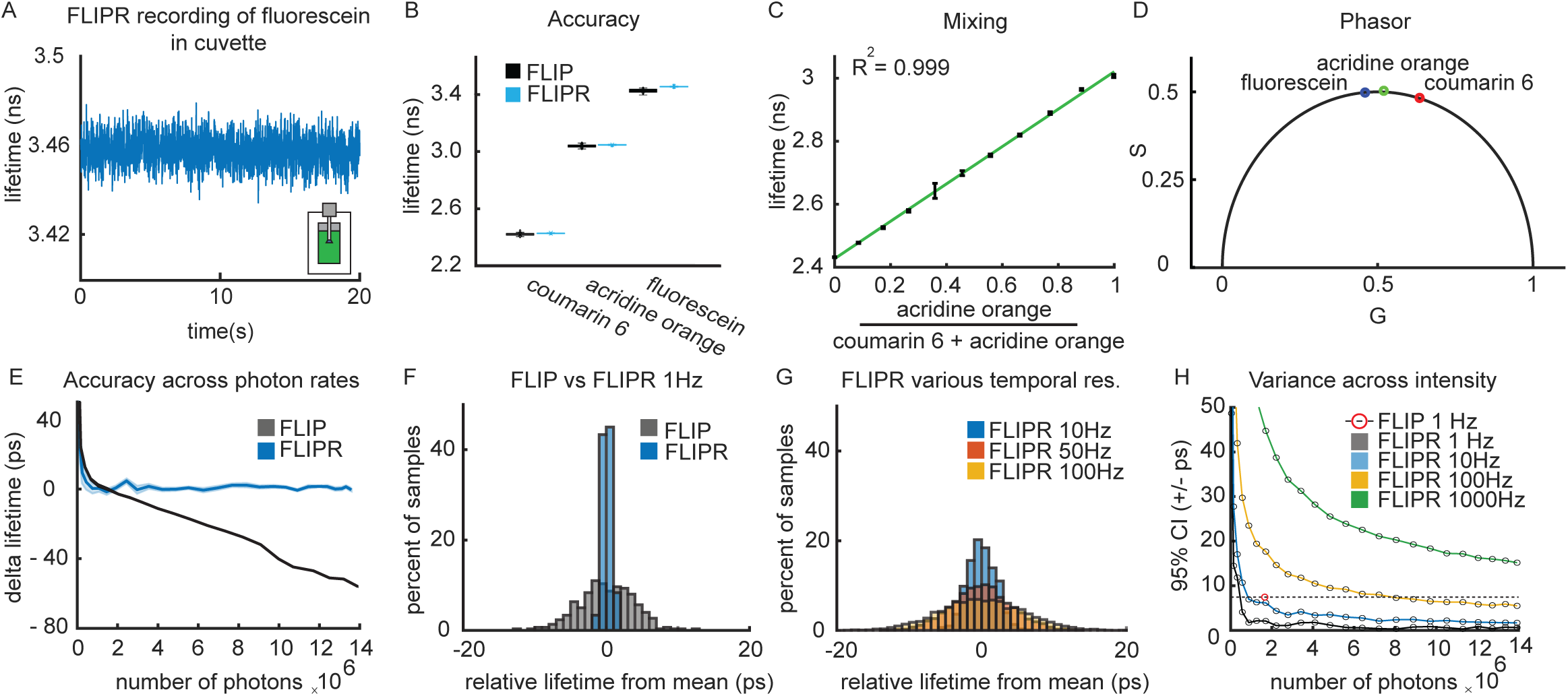
*In vitro* validation of FLIPR **A**, Example fluorescence lifetime of fluorescein measured through a fiber optic placed in a cuvette with FLIPR at 100 Hz. **B**, Comparison of mean fluorescence lifetime estimates of coumarin 6, acridine orange and fluorescein using FLIP (1 Hz) and FLIPR (10 Hz) measured as in panel A. Error bars, 95% confidence interval. **C**, Fluorescence lifetime of different ratios of acridine orange and coumarin 6 to determine linear accuracy of FLIPR measurements. Green line: linear fit, R^2^=0.999. Error bars, SEM **D**, Phasor analysis of FLIPR measurement of coumarin 6, acridine orange and fluorescein, from data displayed in panel B. Accuracy compared to predicted point on the phasor plot was 99.65% SD = 0.27% for G and 99.66% SD = 0.27% S components. **E**, Fluorescence lifetime of coumarin 6 was measured using FLIP and FLIPR with varying number of detected photons. As expected, due to photon pileup, FLIP increasingly underestimates the lifetime at higher photon counts. In contrast, the fluorescence lifetime measured using FLIPR remains stable across a broad range of fluorescence intensities. **F**, Example histogram showing variance of coumarin 6 fluorescence lifetimes measured using FLIP at 1 Hz or FLIPR at 10 kHz followed by sample averaging and down sampling to 1Hz. **G**, Example histograms showing variance of coumarin 6 fluorescence lifetime measured using FLIPR at 10 kHz and down sampled to 10, 50, and 100 Hz. **H**, Variances of coumarin 6 fluorescence lifetimes measured with FLIPR at different temporal resolutions and fluorescence intensities. For comparison the coumarin 6 fluorescence lifetime measured using FLIP at 1 Hz is shown for a power level at which this approach is accurate (red point and dashed line).

GEFI are typically optimized to maximize state-dependent changes in fluorescence intensity. Furthermore, investigators may, depending on experimental needs, perform measurements at different excitation intensities, for example to trade off signal-to-noise of rapid measurements vs. long-term stability. Thus, it is necessary to be able to perform accurate lifetime measurements over a wide range of fluorescence intensities without recalibration of the system.

To examine the stability and accuracy of fluorescence lifetime measurements across intensities, the fluorescence lifetime of coumarin 6 was measured at varying photon rates using FLIP and FLIPR. As expected, lifetimes estimated with FLIP decreased at higher photon rates such that they were only accurate near ∼1 MHz photon detection rate, effectively limiting high precision lifetime measurements to ∼1 Hz. In contrast, FLIPR measurements were stable across the measured range for photon rates (0.5-14 MHz) (**Figure 2E**), permitting lower variance measurements at 1 Hz sampling compared to FLIP (**Figure 2F**). Furthermore, we exploited the ability of FLIPR to operate at high speeds without deadtime to sample the fluorescence lifetime at 10 kHz, allowing us to set final effective temporal resolution by down sampling after acquisition. This permits a user-defined tradeoff between measurement variance and speed (**Figure 2G**) and allows temporal resolutions of 10-1000 Hz while maintaining low variance (**Figure 2H**).

### *In vivo* validation

A key limitation of conventional fiber photometry is its inability to capture absolute signals over long timescales. To examine the capabilities of FLIPR to perform continuous measurements over long timescales *in vivo,* we expressed FLiM-AKAR T391A, a pH stable inactivated protein kinase A fluorescence lifetime sensor, in the Nucleus Accumbens core (NAC) and implanted a fiber optic above the injection site^2^. After 4 weeks of expression, we measured the sensor lifetime daily for 6 days (**Figure 3A-C**) while the mouse freely explored an arena^2^. The fluorescence lifetime of FLiM-AKART391A remained stable across the 2-hour session (**Figure 3B**, mean=1.548 ns, confidence interval (CI)=0.005 ns or 0.33% of the mean). In contrast, the intensity channel showed significant variance in the signal over time, possibly due to motion and hemodynamic artifacts (CI=7.75% of the mean). However, these intensity changes minimally effected the fluorescence lifetime, highlighting the insensitivity of FLIPR to artifacts typically found in intensity fiber photometry (R^2^ = 0.00011). Furthermore, there was no significant change in FLiM-AKAR T391A fluorescence lifetime across days (**Figure 3C**, p = 0.223, one way ANOVA). Similar stability was found for multiple GFP variants whose lifetimes vary over a wide range such as GFP-BrUSLEE, FLiM-AKAR T391A, Superfolder GFP, and GFPivy (**Figure 3D-E**, average lifetimes are 0.678, 1.537, 2.072 and 2.898 ns, respectively) ^2,33–35^ . Lastly, to determine typical variance of *in vivo* fluorescence lifetimes measurements, we measured the fluorescence lifetime of FLiM-AKAR T391A *in vivo* at different excitation powers and calculated the variance of the signal at 10, 100, or 1000 Hz sampling frequencies (**Figure 3F**), demonstrating how users can trade off SNR and long-term stability via selection of sampling frequency and excitation power. For most users, the excellent performance of ∼5 ps CI at 100 Hz measurements at 10 µW excitation power will be adequate.

**Figure 3.**
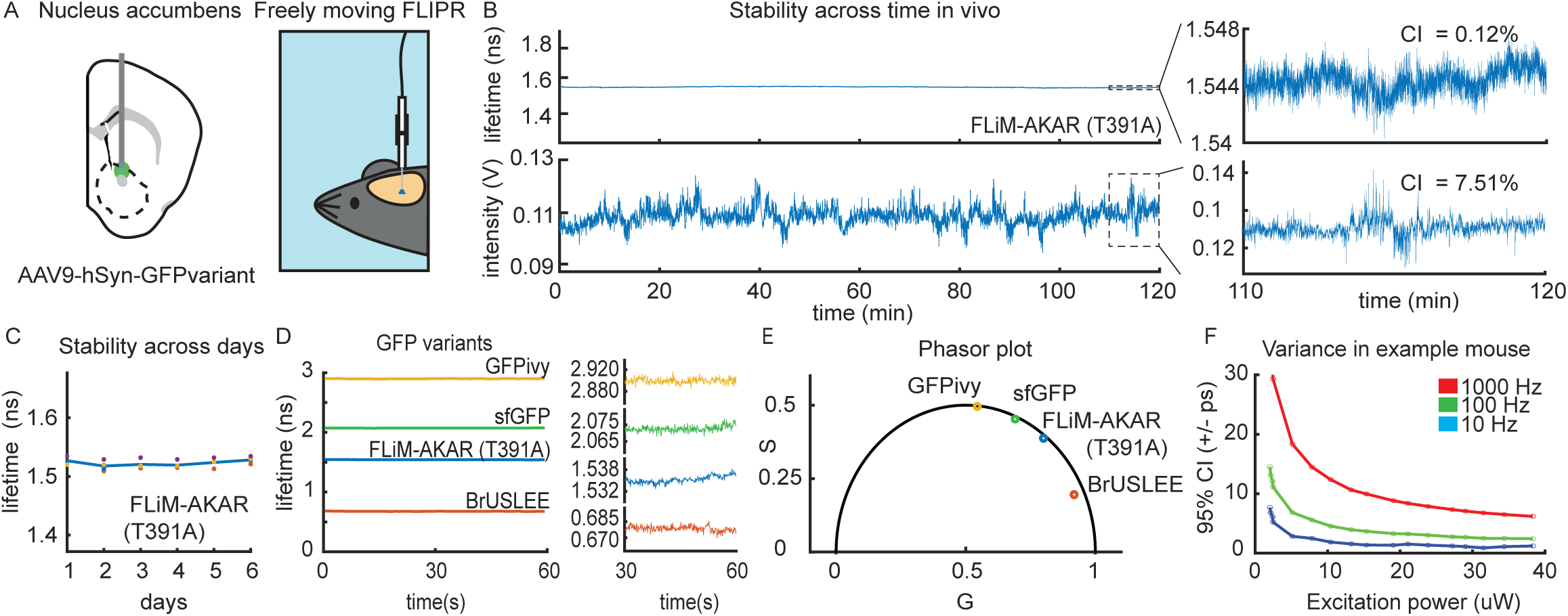
*In vivo* validation of FLIPR **A**, Schematic of experimental setup for *in vivo* validation of the FLIPR system. GFP variants GFPivy, super folder GFP (sfGFP), FLiM-AKAR T391A and GFP BRuSLEE were individually expressed in the NAC. A fiber optic was placed above the injection site and the fluorescence lifetime was measured using FLIPR. **B**, *In vivo* fluorescence lifetime and intensity of FliM-AKAR T391A, a pH stable EGFP containing inactivated PKA sensor, were measured continuously for 2-hours in a freely moving mouse. Fluorescence lifetime was more stable across the session compared to the intensity (intensity CI = 7.75% from mean, fluorescence lifetime CI = 0.35% from mean). Both measurements are shown with the same Y-axis range of mean +/- 20%. On the right are shown data from the last 10 min of recording on an expanded scale with corresponding variance. **C**, Fluorescence lifetime of FliM-AKAR T391A was measured in freely moving mice across 6 days. The fluorescence lifetime did not change across days (p = 0.337, one-way repeated measures ANOVA, n = 4 hemispheres). **D**, Fluorescence lifetimes of GFPivy, sfGFP, FliM-AKAR T391A and BRuSLEE were measured *in vivo* for 60 sec in freely moving mice. Data are shown with all GFP variants on one axis (*left*) to show differences in absolute lifetimes and plotted individually (*right*) to show the stability of each measurement. **E**, Phasor plot analysis of GFPivy, sfGFP, FliM-AKAR T391A and BRuSLEE *in vivo* lifetime measurements. **F**, Variance of FliM-AKAR T391A fluorescence lifetime measured *in vivo* across excitation powers in an example mouse.

### Validation of dLight3.8 to enable absolute dopamine measurement in vivo

We previously uncovered ligand-dependent fluorescence lifetime changes in the new dopamine sensor dLight 3.8 (Roshgadol et al., in preparation). To enable high speed absolute dopamine measurements *in vivo*, we validated the use of dLight3.8 using FLIPR. First, we injected AAV9-cag-dLight3.8 into the Nucleus Accumbens and placed a fiber above the injection site to measure fluorescence lifetime and intensity in freely moving mice (**Figure 4AB**). Fluorescence lifetime was strongly correlated with intensity (**Figure 4B**, R^2^=0.936) during endogenous fluctuations and could be increased or decreased with optogenetic stimulation or inhibition, respectively, of midbrain dopamine (**Figure S3**). The sublinearity of the relationship between lifetime and intensity is expected from the increased brightness of the dopamine bound relative to the unbound sensor and does not reflect electronic artifacts (see methods). To further validate that the fluorescence lifetime reflects the dopamine concentration, we mutated the dopamine binding site of dLight3.8 (D127A) to generate dLight3.8mut (dLight3.8 D127A, **Figure 4C**). dLight 3.8mut had significantly reduced fluorescence lifetime baseline and variance (**Figure 4D-F**). Finally, intraperitoneal (I.P.) injection of SCH23390 (10 mg/kg), a non-competitive antagonist of Type-1 dopamine receptor (D1R), on which dLight3.8 is based, reduced the baseline and variance of dLight3.8 lifetime significantly (**Figure 4G-I**). Although D1R antagonist injection did not affect dLight 3.8mut variance it did slightly increase the lifetime of dLight3.8mut consistent with potential remaining allosteric effects of the compound on the mutated receptor.

**Figure 4.**
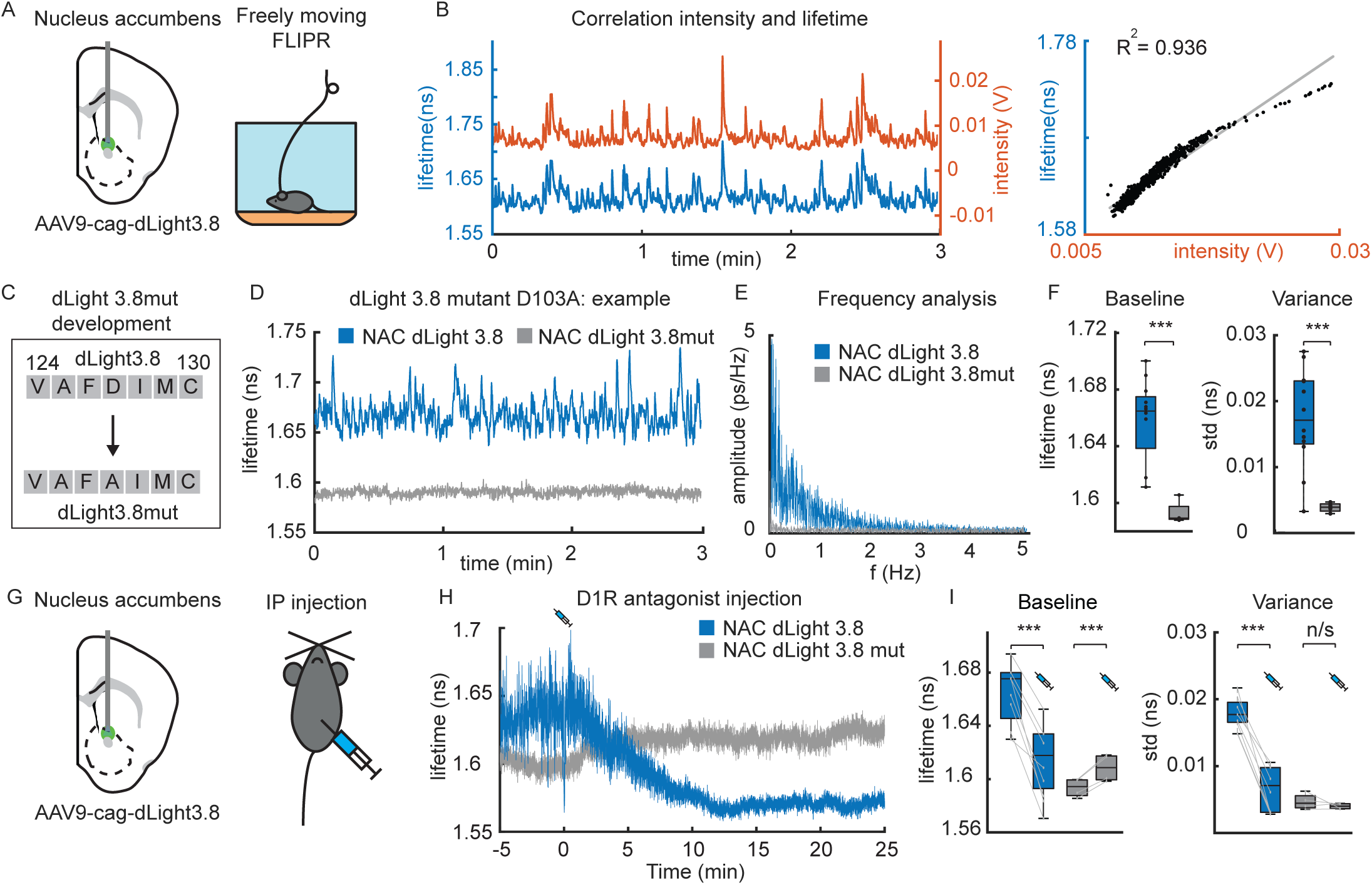
dLight 3.8 is a lifetime reporter of dopamine **A**, Schematic of experimental setup for *in vivo* FLIPR measurements of dLight3.8 fluorescence intensity and lifetime. The dopamine sensor dLight 3.8 was expressed in the NAC and a fiber optic was placed above the injection site. **B**, dLight3.8 fluorescence intensity (red) and lifetime (blue) changes measured *in vivo* correlate (Ieft) nearly linearly (*right*) fit (linear fit is shown, R^2^= 0.936). The sublinear relationship between intensity and lifetime is expected, as the change in intensity ratio between bound and unbound dLight 3.8 due to dopamine binding diminishes with increased dLight3.8 occupancy. **C**, A ligand binding site mutant of dLight3.8 (dLight3.8mut) was generated by changing aspartate (D) 127 to alanine (A). Dlight3.8 and dLight3.8mut were expressed in the NAC for FLIPR analysis in freely moving mice. **D**, Example traces of fluorescence lifetime measurements of dLight 3.8 (blue) and dLight 3.8mut (grey). **E**, Fourier transform frequency analysis of dLight3.8 and dLight3.8mut baseline recordings. **F**, Baseline fluorescence lifetime (*left*) and lifetime variance (*right*) of dLight3.8mut (n=4 hemispheres) are significant reduced compared to those of dLight3.8 (n=10) (two-sample T-test, p=0.0048 and 0.0034, respectively). Error bars: SEM. **G**, Schematic of the experiments used to measure the effects of D1R antagonism on dLight3.8 and dLight3.8mut fluorescence lifetimes. Mice were injected IP with the D1R antagonist SCH23390 (10 mg/kg). **H**, Example traces of *in vivo* fluorescence lifetime measurements of dLight3.8 (blue) and dLight3.8mut (grey) following D1R antagonist injection (t=0 min). **I**, Average mean (*left*) and variance (*right*) of absolute fluorescence lifetimes before and after D1R antagonist injection (lines connect data for individual mice). D1R antagonist injection significantly reduced the lifetime of dLight3.8 (paired sample T-test, p=2.76E-6, n=8) and increased it for dLight3.8mut (paired sample T-test, p=0.0038, n=4). In the same measurements, D1R antagonist decreased the variance of fluorescence lifetime of dLight3.8 (paired sample T-test, p=5.11E-8, n=8) but did not affect that of dLight3.8mut (paired sample T-test, p=0.39, n=4). Baseline values were calculated at t=-5 to 0 min and compared to t=20-25min. Error bars: SEM.

### *In vivo* measurement of absolute tonic and phasic dopamine signals with FLIPR

We examined the ability of measurements of dLight3.8 lifetime with FLIPR to report absolute phasic dopamine signals, detect within-session tonic changes in dopamine, and compare basal levels of dopamine across manipulations and striatal regions. The striatum has functionally distinct regions: the NAC receives dopaminergic inputs from the ventral tegmental area that encode reward prediction errors, whereas the tail-of-striatum (TOS) receives dopaminergic inputs from the substantia nigra pars lateralis that encode threat and salience^36,37^.

We injected dLight 3.8 or dLight3.8mut into the TOS or NAC and placed a fiber optic above the injection site (**Figure 5A-B**). Mice were given food pellets and subject to foot shocks (**Figure 5C-E**). Pellet consumption evoked phasic increases in dopamine in the NAC and TOS, with those in the former being approximately 2-fold larger in magnitude. Foot shocks minimally altered dopamine in the NAC but evoked large phasic dopamine transients followed by a small dip in the TOS. We compared absolute changes in phasic dopamine for NAC food pellet and TOS foot shock response down sampled to 1 Hz vs 20 Hz to highlight the importance of the FLIPR improved sampling rate to accurately estimate the amplitude of phasic transients (**Figure 5F**-**G**). Down sampling distorted the dLight3.8 waveform, underestimating the magnitude of the evoked change in lifetime, and minimizing the differences between signals in the TOS and NAC.

**Figure 5.**
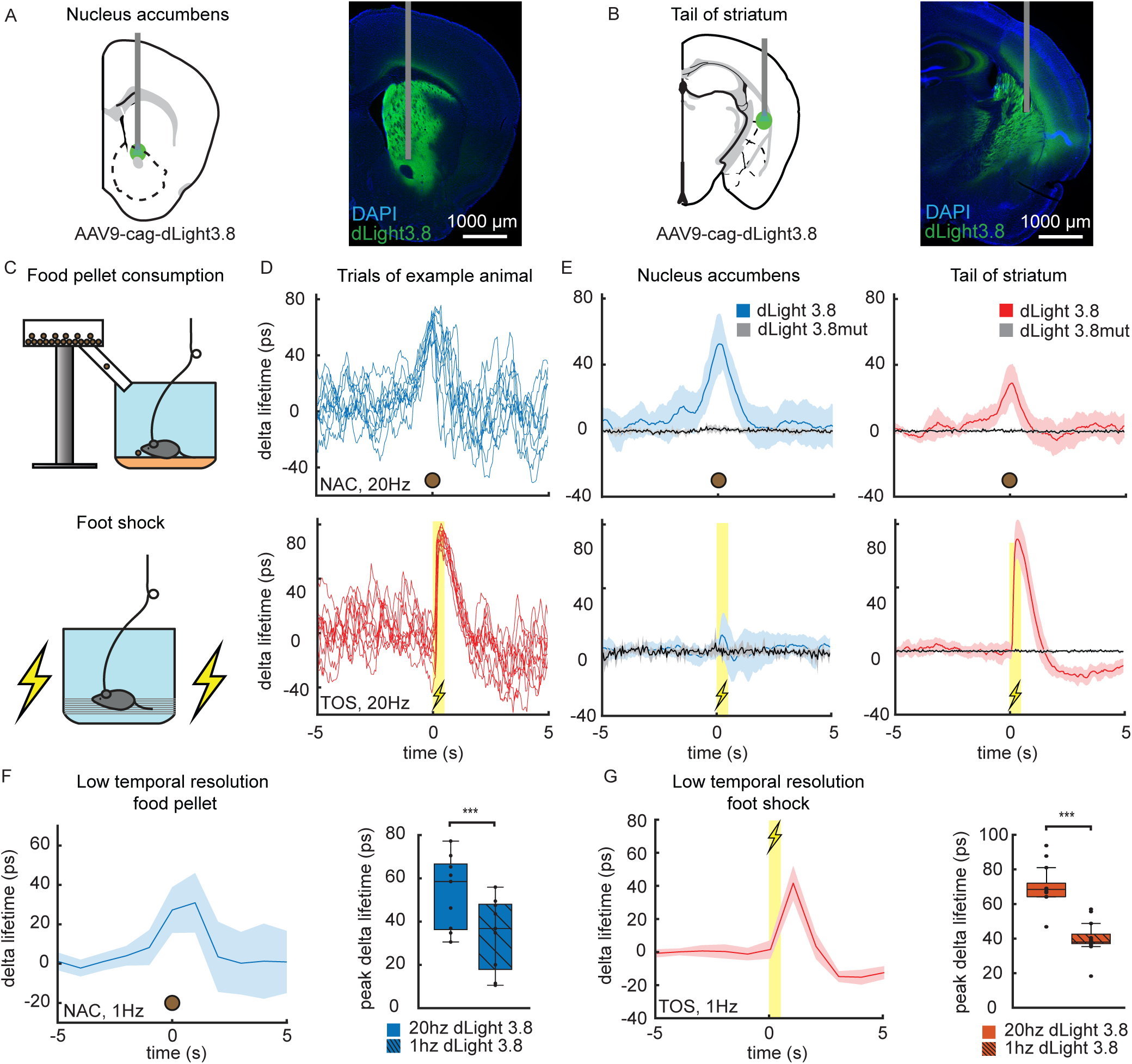
Phasic dopamine in the NAC and TOS across hemispheres to reward and punishment **A**, Schematic of injection of AAV9-cag-dLight3.8 and fiber implantation in the NAC (left) and example histology of fiber placement (right). **B**, Schematic of injection of AAV9-cag-dLight3.8 and fiber implantation in the TOS (left) and example histology of fiber placement (right). **C**, Schematics of devices for spontaneous delivery of food pellets (*top*) or foot shocks (*bottom*) to animals undergoing FLIPR measurements in the TOS or NAC. **D**, Example fluorescence lifetime measurements of dLight3.8 in individual trials (thin lines, t=0 indicates the start of food pellet consumption) collected in NAC (blue, *top*) and TOS (red, *bottom*). **E**, Average of fluorescence lifetime transients of dLight3.8 and dLight3.8mut collect in the NAC (*left*) and TOS (*right*) evoked by food pellet consumption (*top*) (NAC: blue, n=9 dLight3.8, grey n=4 dLight3.8mut; TOS: red, n=8, grey n=4) or foot shocks (bottom) (NAC: blue, n=12 dLight3.8, grey=4 dLight3.8mut; TOS: red, n=13 dLight3.8, grey=6 dLight3.8mut). Traces show the means +/- SEM across animals and hemispheres. **F**, average fluorescence lifetime transient of dLight3.8 in the NACC in response to food pellet consumption, as in panel E (left, top), but down sampled to 1hz (left). NAC dLight3.8 peak delta lifetime was significantly reduced by down sampling during food pellet consumption (right, paired sample T-test, p = 2.1365E-5, n = 9). **G**, average fluorescence lifetime transient of dLight3.8 in the TOS in response to foot shock, as in panel E (right, bottom), but down sampled to 1hz (left). TOS dLight3.8 peak delta lifetime after foot shock was significantly reduced by down sampling (right, paired sample T-test, p = 4.673E-13, n=13).

To compare baseline dopamine in these distinct regions, we measured the fluorescence lifetime of dlight3.8 in the NAC and TOS in freely moving mice in a neutral experimental chamber (**Figure 6A**). Tonic dopamine levels, as reported by dLight3.8 baseline lifetime, were significantly higher in the TOS compared to the NAC (**Figure 6B**). To control for potential contributions of brain autofluorescence to the average lifetime, we investigated the level of autofluorescence *in vivo* across time in the TOS and NAC (**Figure S4**). Autofluorescence is higher in the TOS and has a fluorescence lifetime further from dLight3.8 than in the NAC. However, autofluorescence correction (see methods) maintained the difference in dLight3.8 lifetimes in TOS and NAC and dLight3.8mut lifetimes showed no significant differences between the NAC and TOS (**Figure 6B**), supporting the existence of true differences in tonic dopamine levels in TOS and NAC.

**Figure 6.**
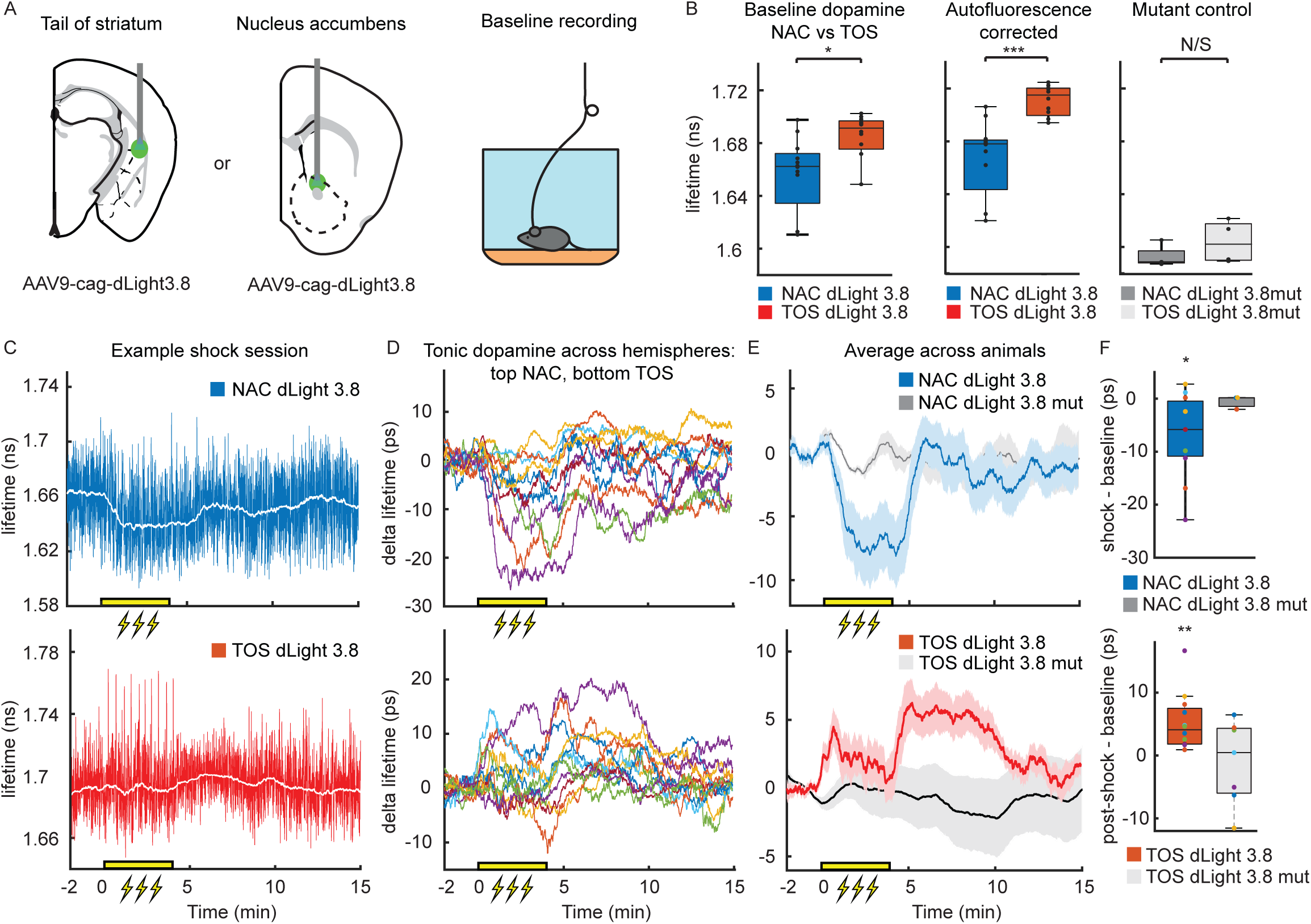
Regional and contextual variation in tonic dopamine measured with FLIPR **A**, Comparison of tonic and phasic DA signals in NAC (*right*) and TOS (*left*) measured with dLight3.8 and FLIPR in freely moving mice. **B**, Basal dLight 3.8 fluorescence lifetime is significantly elevated in the TOS (red, n=12 hemispheres) compared to NAC (blue, n=12) with (*middle,* two-sample T-test, p=3.533e-4) and without (*left,* p=0.0219, two-sample T-test) autofluorescence correction. There is no significant difference of dLight3.8mut fluorescence lifetime between the TOS and NAC (*right*, p=0.264, two-sample T-test, NAC n=4 and TOS n=4 hemispheres). Error bars: 95% confidence interval. The bottom and top of the box are the 25th and 75th percentiles. **C**, Example session dLight3.8 measured in the NAC (*top*) and TOS (*bottom*). Blue and red traces are down sampled to 10 Hz. The white line shows median of the data calculated in a 1 min rolling window . Yellow bar indicates the duration of the foot shock session with one shock delivered every 30 seconds for 4 minutes. **D,** As in C, showing data for individual animals and measurement sites. During the foot shock period tonic dopamine decreased in NAC (*top*) and increases in the TOS (*bottom*). The signal increases further in TOS after the end of the foot shock period. **E**, As in D, showing average data across mice. On average baseline fluorescence lifetime of dLight3.8 decreased in NAC during foot shock (*Top*, blue n =11, grey n = 3) and increased in the TOS after foot shock (*Bottom*, red n =11, grey n=7). Traces represent the mean across animals, yellow bar indicates duration of the foot shock session, one 500ms shock every 30 seconds for 4 minutes. Error bars, SEM. **F**, dLight3.8 mean fluorescence lifetime in the NAC was significantly decreased during shock trials (0-5 minutes) relative to baseline (-2 – 0 minutes) (*top*, p = 0.017, two-sample T-test, blue dLight3.8 n = 11; p = 0.489, dark grey dLight3.8mut n = 3). dLight3.8 mean fluorescence lifetime in the TOS was significantly increased after shock trials (5-10 minutes) relative to baseline (-2 – 0 minutes) (*top*, p = 0.002, two-sample T-test, red dLight3.8 n = 11; p = 0.621, light grey dLight3.8mut n = 7). Boxplot error bars are the 95% confidence interval, the bottom and top of the box are the 25th and 75th percentiles.

Finally, to investigate dynamic changes in tonic dopamine across the session, we extracted basal dLight3.8 lifetime by performing a moving median filter (60 seconds) over the entire session of the previously collected foot shock data (**Figure 6C**). As baseline dopamine varies animal to animal, we calculated the delta lifetime through baseline (-2 – 0s) subtraction. Although there were no foot shock-evoked phasic dopamine transients in NAC (**Figure 5E**), we observed a dynamic reduction in tonic dopamine during the 4-min period in which shocks were delivered. Tonic dopamine in NAC returned to the pre-shock baseline with ∼1 min of the conclusion of the uncued shock period (**Figure 6C-F**). In contrast, TOS tonic dopamine increased slightly during the shock period and increased for ∼5 min after the shock period concluded before returning to baseline (**Figure 6C-F**). Similar changes in fluorescence lifetime were not observed in NAC or TOS in mice expressing dLight3.8mut (**Figure 6**). Thus, FLIPR, used in conjunction with dLight3.8, can concurrently detect fast and slow changes in dopamine in the striatum, revealing regional specialization of tonic and phasic signaling evoked by behaviorally relevant events.

## Discussion

We designed and used FLIPR, a new frequency-domain fluorescence lifetime photometry system that enables high-speed fluorescence lifetime measurements in the brain of freely moving mice. FLIPR relies on an optimized analog-computing unit for high speed (1-1000 Hz), high resolution and accurate fluorescence lifetime measurements. FLIPR is more affordable and robust than previous time domain FLIP systems and is built from simple and low-cost analog components, making measurement of fluorescence lifetime *in vivo* accessible to the research community. Here we demonstrate the capabilities of FLIPR by exploiting the dopamine-dependent changes in fluorescence lifetime of the new dopamine sensor dLight3.8. FLIPR reveals heterogeneity in both phasic and tonic dopamine signaling across striatal regions and behavioral contexts. In short, FLIPR facilitates the adoption of robust fluorescence lifetime photometry, allowing the community to take advantage of a new wave of fluorescence lifetime reporters being generated.

### Comparison to other approaches for lifetime photometry

A key technical advantage that lowers the cost and complexity of FLIPR is that, because the computation is done in an analog processing unit, the sampling frequency by the digitizer can occur at the sampling frequency that the user desires, not at the high rates (∼10 GHz) necessary to measure single photon arrival times. This is in contrast with most previous fluorescence lifetime photometry systems, which rely on time correlated single photon counting systems (TCSPC) that measure arrival times of individual photons relative to a sync pulse with picosecond (ps) resolution. Because of the complex electronics and timing mechanisms, these time counting systems also typically have photon detection “deadtimes”, preventing counting of photons up to ∼100 ns after a photon is processed. This caps the theoretical maximal photon counting rates of the system. In practice, other technical concerns such as photon pileup reduce the maximal detection rate to ∼1 MHz which, when coupled with the need to collect several 100,000s of photons to provide an accurate lifetime measure, limits the effective temporal resolution of lifetime measurements to ∼1 Hz ^9,17,38^. Photon rates beyond the limits imposed by electronics artificially lowers the lifetime estimate, introducing signal-dependent and dynamic measurement errors that can be difficult to detect and avoid *a priori*. There is no theoretical or technical limit on lifetime measurements at slow time scales (i.e., to monitor baseline changes) but due to limitations described above it is not possible to measure fast and slow signals with the same system.

Frequency domain lifetime measurement systems do not have photon detection deadtimes and every photon that is detected in the PMT contributes to the calculation of the fluorescence lifetime^20^. This, and cost savings, motivated the development of the original analog processing unit for fluorescence lifetime measurements^20^. To achieve accurate, fast, and high dynamic range measurements with FLIPR, we suppressed electronic reflections to reduce noise, fixed phase delays, and properly power balanced signals across the system. These improvements made FLIPR accurate across a large range of detected photon rates (0.5-14 MHz), allowing for lifetime measurements at 1-1000 Hz with ps variance. Although properly used time counting systems are the gold-standard for lifetime measurements, we show that, at the same rate of photon detection by the PMT, FLIPR has lower measurement variance, likely due to the rejection of photons by the electronics in time domain systems.

The ability of FLIPR to accurately measure lifetime across a broad range of light levels, permits its use for the analysis of fluorescence transients from sensors that have large conformation-dependent intensity changes in addition to lifetime changes. We demonstrate experimentally that, as is for a true measure of the changes in lifetime of individual sensor molecules, FLIPR measurements are insensitive to many artifacts and sources of variance that complicate comparison of photometry signals across time, sessions, and animals^12^. Lastly, extracting fluorescence lifetime from the analog outputs of FLIPR is straightforward requiring only simple matrix calculations^20^. Thus, FLIPR will be easy to implement for labs with fiber photometry experience, as surgical and behavioral requirements are identical for both techniques.

### Comparison to other approach for absolute measurements of analytes *in vivo*

Fast scan cyclic voltammetry (FSCV) is a powerful approach to directly measure compounds that can undergo ox/redox reactions at the surface of a carbon microfiber. FSCV is typically challenging to implement *in vivo* due to the complexity of data collection and analysis, caused by factors such as interference from other molecules, degradation of the electrode, and hemodynamic, electrical and motion artifacts ^39–42^. Long-term chronic FSCV measurements are particularly difficult *in vivo* due to the prolonged degradation of the carbon fiber working electrode and reference electrode, which complicates background subtraction and voltammogram identification^43^. However, calibration of a FSCV probe *ex vivo* before or after implantation does allow for absolute concentration detection^39^. In addition, a recent preprint proposed a new method by which FSCV can be used in a ratiometric mode to derive relative phasic and tonic dopamine levels ^44^.

FLiPR has several advantages over fast scan cyclic voltammetry (FSCV), such as ease of adoption by laboratories, the ease of use in freely moving animals, the number of different molecules and processes that can be measured, the ability to measure multiple molecules simultaneously, and the stability of absolute measurements across days using the same probe. FLIPR has the same surgical and behavioral requirements as fiber photometry and does not suffer from probe degradation, allowing for absolute measurement of molecules for the duration of sensor expression, typically for multiple months.

Although microdialysis allows for the capture of the absolute concentration of many molecules, microdialysis is typically slow due to high sampling volumes (sampling rate of minutes), may not accurately report concentration due to rapid molecule degradation and depletion, causes acute damage during measurement and is not compatible with real time measurements of neuronal signals^45^.

### Tonic and phasic dopamine signaling

Tonic and phasic dopamine release determine the activation of dopamine receptors and subsequent downstream intracellular signaling, which govern neuronal plasticity and excitability^46,47^. Due to technical limitations, most experimental efforts have focused on understanding phasic dopamine with a few efforts examining tonic signaling ^10^. However, to truly understand dopamine function, it is crucial to measure phasic and tonic dopamine simultaneously, as the downstream impacts of phasic transients depend on baseline receptor occupancy, which is set by tonic dopamine levels^32,48^. The interplay between tonic and phasic dopamine has been theorized to determine learning rates and to contribute the cause or treatment of several neuropsychiatric disorders, including Parkinson’s disease, drug addiction, schizophrenia and attention deficit hyperactivity disorder^49–53^. The ability to measure both phasic and tonic dopamine simultaneously using FLIPR allows for in-depth analysis of the causes of these dysfunctions in disease models and for improved *in vivo* screening of candidate therapeutics.

We applied FLIPR to investigate tonic and phasic dopamine signaling in two regions of the striatum, the NAC and TOS, in response to delivery of appetitive and aversive stimuli. We observed increases in phasic dopamine during food pellet consumption in the NAC and to foot shocks in the TOS, as expected based on the known functions of these regions, respectively, in reward and threat processing^36,54^. We show that fast acquisition is necessary to accurately capture the timing and magnitude of phasic dopamine and compare these signals across regions and conditions.

Foot shocks do not evoke phasic dopamine transients NAC, which may lead to the conclusion that foot shocks do not affect NAC dopaminergic circuity. However, FLIPR reveals that NAC basal dopamine decreases during the period that foot shocks are delivered and that, in contrast, tonic dopamine in the TOS increases after the shock period ends. Such changes in baseline may facilitate or repress phasic dopamine transients and the intracellular signaling cascades that they activate. Therefore, decreased tonic dopamine in the NAC may alter reward value and reward-based learning during periods in which the animal detects threats. Similarly, the transient increase in tonic dopamine after the shock period in the TOS may alter threat sensitivity.

Highlighting the ability to compare across regions and animals, we also examined absolute baseline dopamine in the TOS and NAC. We observed higher levels of tonic dopamine in the TOS compared to the NAC. The biological reasons for differences in tonic dopamine may reflect differences in the properties of dopamine axons in each region or the effects of local signals that impact dopamine release and clearance.

Altogether, FLIPR reveals that phasic and tonic dopamine differentially and dynamically change in a brain region and context specific manner. Given the ease of tonic dopamine detection using FLIPR, we expect that further richness in the dynamic regulation of tonic dopamine will be uncovered.

### Limitations and future

A limitation of lifetime photometry is the current relative lack of fluorescence lifetime sensors compared to intensity-based ones. Lifetime changes are unexpected in sensors that couple biological processes to allosteric changes that cause rearrangements of circularly permuted GFP (cpGFP). Nevertheless, lifetime changes have been observed in such sensors, including in dLight3.8 and other similar sensors^18^. Discovery that other pre-existing sensors can operate in lifetime mode as well as concerted efforts to develop new lifetime fluorescence sensors will likely greatly expand the cellular and circuit processes that can be monitored with FLIPR. Sensors designed specifically for lifetime measurements ideally have only small intensity changes, which facilitates accurate measurement of lifetimes in the dim state and increases the linearity of lifetime changes relative to the biological process of interest.

A second current limitation is that, although FLIPR provides an absolute measurement that can be compared across time, brain regions, and animals, it is not a calibrated measure. In theory, fluorescence lifetime measurements obtained with FLIPR can be calibrated to different concentrations of molecules in the same manner as FSCV; however, absolute fluorescence lifetimes of sensors *in vitro* and *in vivo* may differ. Thus, translating the absolute lifetime measurements provided by FLIPR into concentrations of analytes is still difficult.

The FLIPR system described here was built by hand and is not difficult to reproduce. However, it does require attention to detail and may be challenging for some. Production of a printed circuit board that can be ordered with surface-mount electronics in place will simplify adoption. In addition, replacement of PMTs with silicon-based detectors and the pulsed laser with other high frequency (> 20 MHz) modulated light sources may simplify power control and permit easily changing the excitation and emission spectra to accommodate new fluorophores. This would also simplify the development of multi-color FLIPR systems, which could be used to measure multiple molecules simultaneously. Despite these limitations, FLIPR allows for absolute fluorescence lifetime measurement at low cost, permitting fast and slow neuronal signals to be measured and compared across time scales, animals and conditions in an accessible manner.

## Methods

### Animals

Experimental manipulations were performed in accordance with protocols approved by the Harvard Standing Committee on Animal Care following guidelines described in the US National Institutes of Health Guide for the Care and Use of Laboratory Animals. *C57BL/6J* (000664) mice and *DAT-IRES-Cre* (B6.SJL-Slc6a3tm1.1(cre)Bkmn/J, 006660) were acquired from the Jackson Laboratory. Mouse ages were typically 2-4 months, and male and female mice were used in approximately equal proportion. Mice were housed on a 12hr dark/12hr light reversed cycle. As measurements performed here have never been performed before, no power-based sample size calculation could be performed.

### Viruses

Recombinant adeno-associated viruses (AAVs) of serotype 9 or DJ/9 were used to express transgenes of interest in either Cre-recombinase dependent or independent manner. AAVs were packaged by commercial vector core facilities (Janelia Vector Core and UNC Vector Core) and stored at −80°C upon arrival. Viruses were used at a working concentration of 10^12^ to 10^14^ genomic copies per ml. 250-300 nl of virus was used for all experiments. The following viral plasmids are available on Addgene: AAV-FLEX-FLIM-AKART391A (#60446) and AAVDJ9-nEF-Con/Foff 2.0-ChRmine-oScarlet (#137161). AAV-hsyn-dLight3.8mut, AAV-hsyn-sfGFP, AAV-hsyn-BrUSLEE and AAV-hsyn-Ivy are available upon request to Dr. Bernardo Sabatini. AAV-cag-dLight3.8 is available at UNC neurotools.

### Surgery

Mice were given *ad libitum* oral carprofen one day before surgery. During the stereotactic surgery, inhaled isoflurane was used as anesthesia. Surgeries were performed using a stereotaxic frame (David Kopf Instruments) and 250-300 nl of virus was injected stereotactically at the following titers: AAV9-cag-dLight3.8 1 x 10^13^gc/ml, AAV9-hsyn-dLight3.8mut 2.1 x 10^13^ gc/ml, AAV9-hsyn-sfGFP 1.4 x 10^12^ gc/ml, AAV9-hsyn-BrUSLEE 5.4 x 10^12^ gc/ml, AAV-hsyn-Ivy 7.8 x 10^12^ gc/ml, AAV-FLEX-FLIM-AKART391A 7.6 x10^12^, AAVDJ9-nEF-Con/Foff 2.0-ChRmine-oScarlet 8.7 x10^12^ gc/ml and AAV9-CAG-FLOXed-stGtACR2-FusionRed 7.6 x10^12^ gc/ml. The following coordinates were used (anteroposterior (AP), medio-lateral (ML) relative from bregma; dorsoventral from brain surface):

NAC: +1.2 AP, +/−1.3 ML, -4.1 DV

Nacl: 1.55 AP, +/-1.4 ML, 4.25 DV

VTA: -3.135 AP, +/−0.5 ML, -4.4 DV

TOS: -1.4 AP, +/- 3.28 ML, -2.45 DV

SNPL: -3.1AP, +/- 2.1 ML, -3.6 DV

All coordinates are in millimeters. For fluorescence lifetime photometry or optogenetic experiments, an optical fiber (MFC_200/230-0.37_4.5mm_MF1.25_FLT mono fiber optic cannula, Doric Lenses) or MFC_200/230-0.37_3mm_MF1.25 was implanted 100-200 μm above the injection site.

### Behavior

For all FLIPR measurements, mice were connected to a patch cord that attached to the FLIPR system as in Figure 1. For fluorescence lifetime baseline recordings, mice were placed in an 8x8 inch black acrylic box. For foot shock experiments, mice were placed in a white plexiglass box with a metal barred floor (ENV-005A, Med Associates ) via which foot shocks can be delivered. Mice received 10-foot shocks (0.5 mA, 500 ms) per session. The foot shock delivery was controlled through a programmable microcontroller (Arduino Uno, Arduino) running a custom script. For food reward experiments, mice were food restricted such that they remained at 80∼95% of their initial weight. Mice were given 2-3 g of regular chow daily in addition to the variable number of 20 mg dustless precision chocolate flavor pellets (F05301, Bio Serv). Animals were habituated to an 8x8 inch black acrylic box for 5 – 10 min and received a pellet every minute from an automatic pellet dispenser (ENV-203-20, Med Associates) for a total of 10 pellets. Animal movements were captured by a camera (FL3-U3-13E4M, PointGrey). Bonsai 2.5.1 software controlled the pellet dispenser and was synchronized with the FLIPR MATLAB 2021 software.

### Histology

Mice were anesthetized with isofluorane inhalation. Mice were perfused transcranially with PBS followed by 4% PFA in PBS. Brains were left overnight in 4% PFA in PBS and then transferred to PBS. Brains were sliced (50-70 μm thickness) using a vibrating blade microtome (Leica Biosystems VT1000S), were mounted on glass slides using ProLong Diamond Antifade Mountant with DAPI (ThermoFisher Scientific), and imaged using an Olympus VS200 slide scanning microscope.

### Pharmacology

D1R antagonist SCH23390 (Tocris, #0925) was diluted in sterile saline and intraperitoneally (IP) injected at 1 mg/ml at 10 mg/kg using a fine syringe (BD 324911, BD Bioscience). Typically, injections were completed 10-60 s after the start of scruffing.

### Fluorescence lifetime photometry

FLIP was performed using the apparatus shown in Figure S1. A 473 nm pulsed laser (BDS-SM-473-FBC, Becker & Hickl) produced a 50 MHz pulsed laser beam that passed through a Thorlabs polarizer (WPH05ME-488, Thorlabs), and was modulated using an AOM (MTS110-A3-VIS, AA optoelectronic) for power control. AOM attenuation was adjusted by an analog output module (NI-9261, National Instruments) controlled by MATLAB software. The laser beam was centered using a pair of visible coated mirrors (BB1-E01 and KCB1, Thorlabs) and passed through a rotary neutral density filter (NDC-50C-4M-A, Thorlabs) for manual power modulation. The beam reflected on a 488 nm dichroic (Di02-R488-25x36, Semrock) mounted in DFM1-P01, Thorlabs) and was launched into a patch-cord (MFP_200/220/900-0.37_2m_FCM-MF1.25(F)_LAF, Doric) by an additional mirror and an achromatic doublet lens (AC254-050-A-ML, Thorlabs). Emission light was collected using the same fiber implant and patch cord. Emission light was directed through the same lens, mirror (M3) and dichroic. Emission light was filtered through an interference filter (FF01-525/35-25, Idex HS). Thorlabs kinetic magnetic cube (DFM1-P01, Thorlabs) containing a mirror (MGP01-350-700-25x36, Semrock) was used to direct the emission light into a hybrid PMT (HPM-100-07-Cooled, BH) controlled by DCC-100-PCI (BH) for FLiP. An empty Thorlabs kinetic magnetic cube was used to direct the emission light into a GaAsp PMT (PMT2101, Thorlabs). All components were light shielded using a variety of Thorlabs lens tubes.

For time domain FLiP, the pulsed laser synchronization port and a hybrid PMT were connected to a time correlated single photon counting system (SPC-830, Becker & Hickl), which detected the time difference between the laser pulse and photon arrival. Data was collected at 1-5 s intervals, with 0.15 s of data transfer time using custom software written in MATLAB 2012a. FLIP data was collected using a rate of 200 kHz–1 MHz accepted photons, except for the data shown in Figure 2E which are shown as a function of variable photon count rates detected by the PMT. Typical excitation power used was 0.5-18 μW. For accuracy comparison experiments, fluorescence lifetime of fluorescent compounds was calculated through single exponential fitting of the photon histogram for each measurement point as previously described ^2,55^. For all other experiments, the average fluorescence lifetime was calculated from the single photon histogram using equation 1:

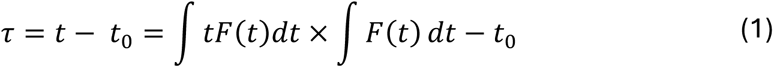

wherein *τ* is the average fluorescence lifetime, t is the average arrival time with respect to the laser pulse, F(t) is the number of photons at time bin t and t_0_ is the timebin corresponding to the peak of the exponentially fitted histogram. The single photon histogram was truncated at 8 – 11.7ns after t_0_ to reject delayed reflection autofluorescence from the system.

For frequency domain FLIPR, the PMT and laser synchronization port was connected to custom made analog phase processing unit (Figure 1A). The laser synchronization pulse was connected to a low pass filter (BLP-70+, Mini-circuits), 20dB attenuator (HAT-20+, Mini-circuits), low noise amplifier (ZX60-P103LN+, Mini-circuits) and 4-way power splitter (ZSC-4-3+, Mini-circuits). From the power splitter each channel was connected to a phase shifter (TB-JSPHS-51+, Mini Circuits), a channel specific delay line (141-## BM+, Mini-circuits), a low noise amplifier (ZX60-P103LN+, Mini-circuits), low pass filter (BLP-70+, Mini-circuits) and LO port of a level 17 frequency mixer (ZX05-1HW-S+, Mini-circuits). The PMT (PMT2101 with internal transimpedance amplifier, Thorlabs) voltage output was connected to a low pass filter (BLP-70+, Mini-circuits), and a bias tee (ZFBT-4R2G+, Mini-circuits), which split the low frequency intensity information (VDC) from the high frequency phase signal (VRF). The VDC was captured using an analog data acquisition module (NI 9215, National Instruments). The VRF signal passed through a low noise amplifier (ZX60- P103LN+, Mini-circuits), and 20dB attenuator (HAT-20+, Mini-circuits). The VRF was connected to a 4-way splitter (ZSC-4-3+, Mini-circuits) and connected to the RF port of frequency mixers. The intermediate frequency port of the frequency mixers was connected to bias tee’s (ZFBT-4R2G+, Mini-circuits). The high frequency port of the bias tees was connected to a 30 dB attenuator (HAT-30+, Mini-circuits) and terminated at 50 Ohm using a resistor. The low frequency port of the bias tee was connected to a 1 KHz low pass filter (EF110, Thorlabs) and a 24-bit analog input module (NI 9239, National instruments). See Table 1 for a detailed list of the components required to build a FLIPR system. Data was collected and processed in real time using custom MATLAB software. Fluorescence lifetime was calculated using equation 2

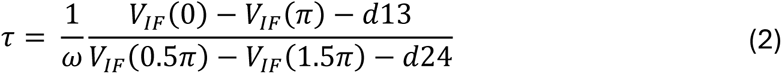

wherein τ is the average fluorescence lifetime, ω is the angular frequency, VIF (0 – 1.5π) are the mixer outputs from phase shifted channel 1-4, d13 and d24 are channel specific offset factors determined during calibration. Data was averaged using a moving mean and down sampled to the required frequencies.

Fluorescence intensity was calculated using equation 3:

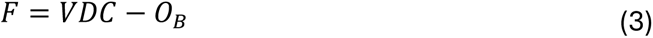

wherein F is intensity signal corrected for baseline voltage, VDC is the voltage coming from the bias tee and O_B_ is the offset of the bias tee determined during calibration.

G and S phasor components were calculated using equation 4 and 5, respectively.

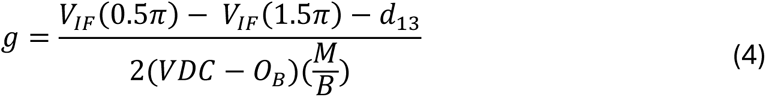

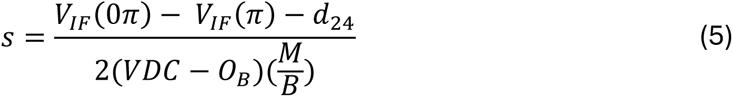

wherein the M is the mixer conversion loss and B is the bias tee conversion loss, measured through calibration. Lifetimes with a single exponential decay have phasor components located on a semi circle termed the universal circle (centered at g = 0.5, s = 0, radius = 0.5), while lifetimes with a multi-exponential decay fall outside of the universal circle. Phasor location can be used to decompose complex mixtures of lifetime agents into individual species, such as the ratio between bound and unbound GEFI or GEFI fluorescence and autofluorescence.

Calibration of the FLIPR system was achieved through the measurement of fluorescence lifetime standard Coumarin 6 (2.5x10^-5^ mg/ml, Sigma Aldrich) dissolved in ethanol which has a known fluorescence lifetime of 2.39-2.41ns. Coumarin 6 solution was measured by FLIPR through a fiber implant of the same length as implanted in animals. Calibration of the FLIPR system was achieved by a calibration protocol described in detail elsewhere^20^.

Two modifications to this protocol were made. First, the polynomial fitting of the phase calibration curve was used to reduce the impact of electrical noise and correctly extract the phase shifter bias voltage. In addition, a second phase calibration was performed after signal path offset and M/B coefficient calibration, to include these factors in phase calibration measurement.

Autofluorescence correction was performed using an intensity weighted average function to extract sensor fluorescence lifetime from measured autofluorescence.

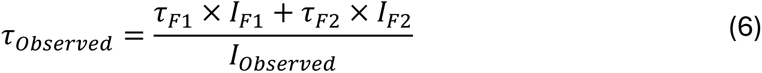

Wherein, τ_Observed_, τ_F1_ and τ_F1_ represent the average, fluorescent source 1 and fluorescent source 2 lifetime respectively, and I_Observed_, I_F1_ and I_F2_ represent the measured, fluorescent source 1 and fluorescent source 2 intensity, respectively. Autofluorescence correction was performed based on empirically determined levels of autofluorescence caused by fiber implantation, tissue damage and viral transfection in the TOS and NAC (S4). Autofluorescence correction was only performed when comparing across brain regions, as autofluorescence contribution errors in delta lifetime calculations were minimal.

### Optogenetic manipulation

To excite ChRmine, a 635 nm laser (Optoengine) was modulated using an AOM at 15 Hz, for 500 ms with 5 ms pulsewidth at 1.1, 1.9 and 2.4 mW. To excite stGtACR2, a blue 488 nm laser (GEM 473, Laser Quantum) was modulated using an AOM for a duration of 5 seconds continuous illumination at 6mW. Light was delivered to the targeted brain region using a 200 µm Doric patch cord. Laser light power was calibrated at the tip of the patch cord using a Thorlabs digital optical power meter (PM400 and S120C).

### Quantification and statistical analysis

Data was analyzed using custom scripts in MATLAB (v2021a and v2024a) and Python (v3.8) and using the data analysis toolbox in excel (v2410). No statistical sample size pre-calculation was performed. Experimenters were not blinded to animal groups, as pilot sensor characterization clearly distinguished dLight 3.8 from dLight3.8mut and fiber placement TOS and NAC. Paired sample T-test, two sample T-test and one-way repeated measures ANOVA statistical tests were applied to data as indicated in figure legends. No multiple comparison correction was performed as none of the multiple comparison tests were significant. In figures, single star (*), double star (**) and triple star (***), represent p < 0.05, p<0.01 and p<0.001, respectively.

## Author contributions

B.L. and B.L.S. designed the FLIPR system. B.L., E.S. B.R. and C.Z. build the FLIPR system. B.L. and S.J. build the FLIP system. B.L., B.L.S., R.A., T.K. and P.R. designed all validation and proof of principle experiments. B.L., T.K. E.S. and M.S. collected all data. J.C. and L.T. developed and provided the dLight3.8 sensor.

## Acknowledgements

We thank all Sabatini lab members for discussions, Yide Zhang for discussing the iFLIM software, Ofer Mazor of the Harvard Medical School Research Instrumentation Core for designing and building the box housing the FLIPR electronics. We thank the Harvard Medical School Machine shop for building components of the behavioral system and patch cord holders required for in vitro characterization. This work is supported by grants to B.L.S. (NIH R35NS137336), B.L. (PhD fellowship, Boehringer Ingelheim Funds) and T.K. (NIH F30DA061675).

## Conflict of interest

Authors declare no conflict of interest.

## Notes

### Competing Interest Statement

The authors have declared no competing interest.

