## Supplementary figures and images for "Absolute measurement of fast and slow neuronal signals with fluorescence lifetime photometry at high temporal resolution"

### Supplemental figures

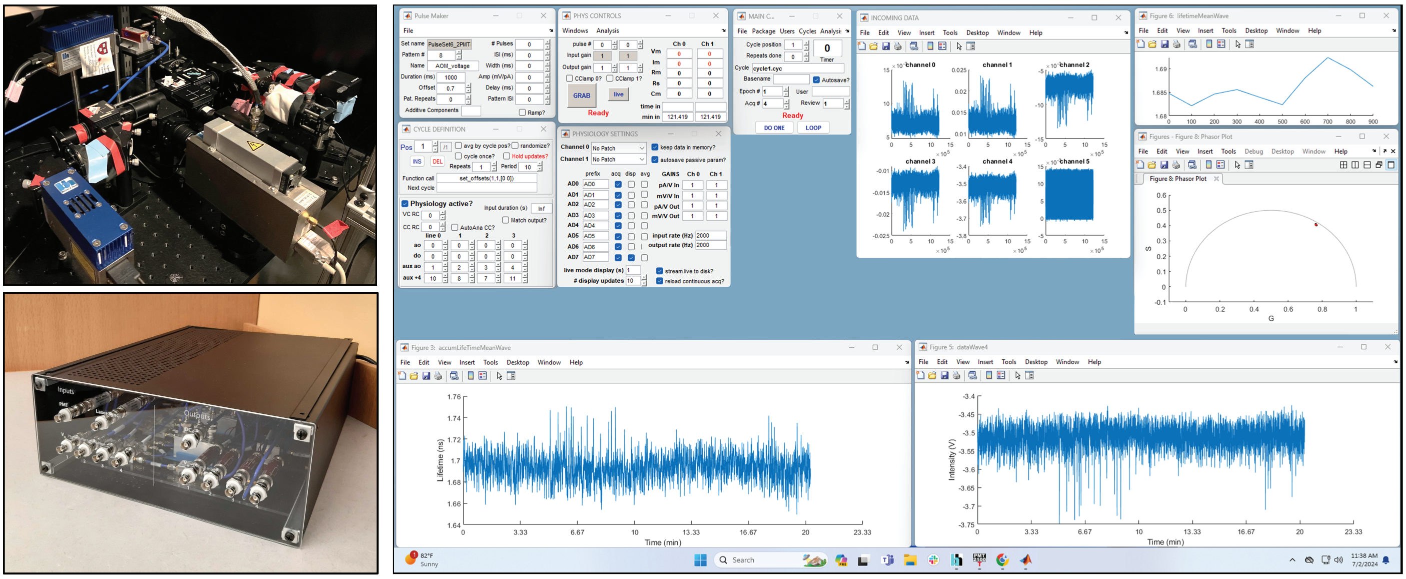

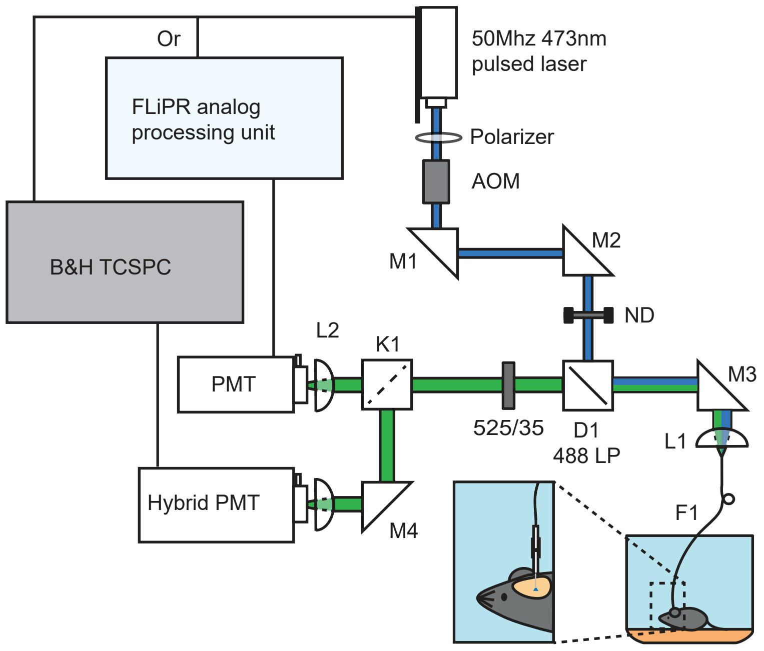

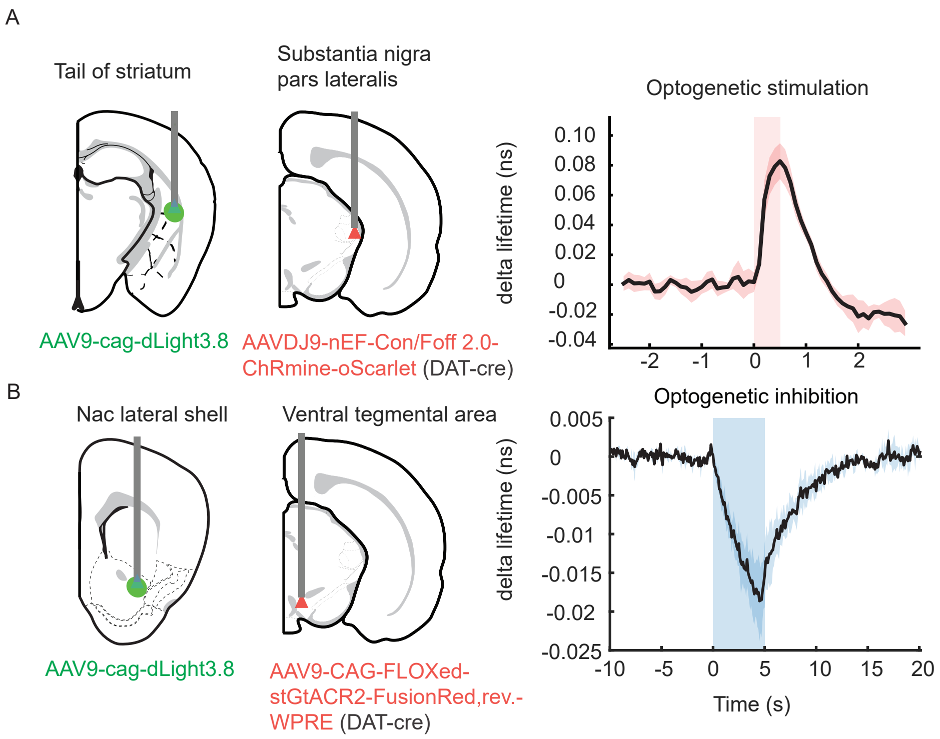

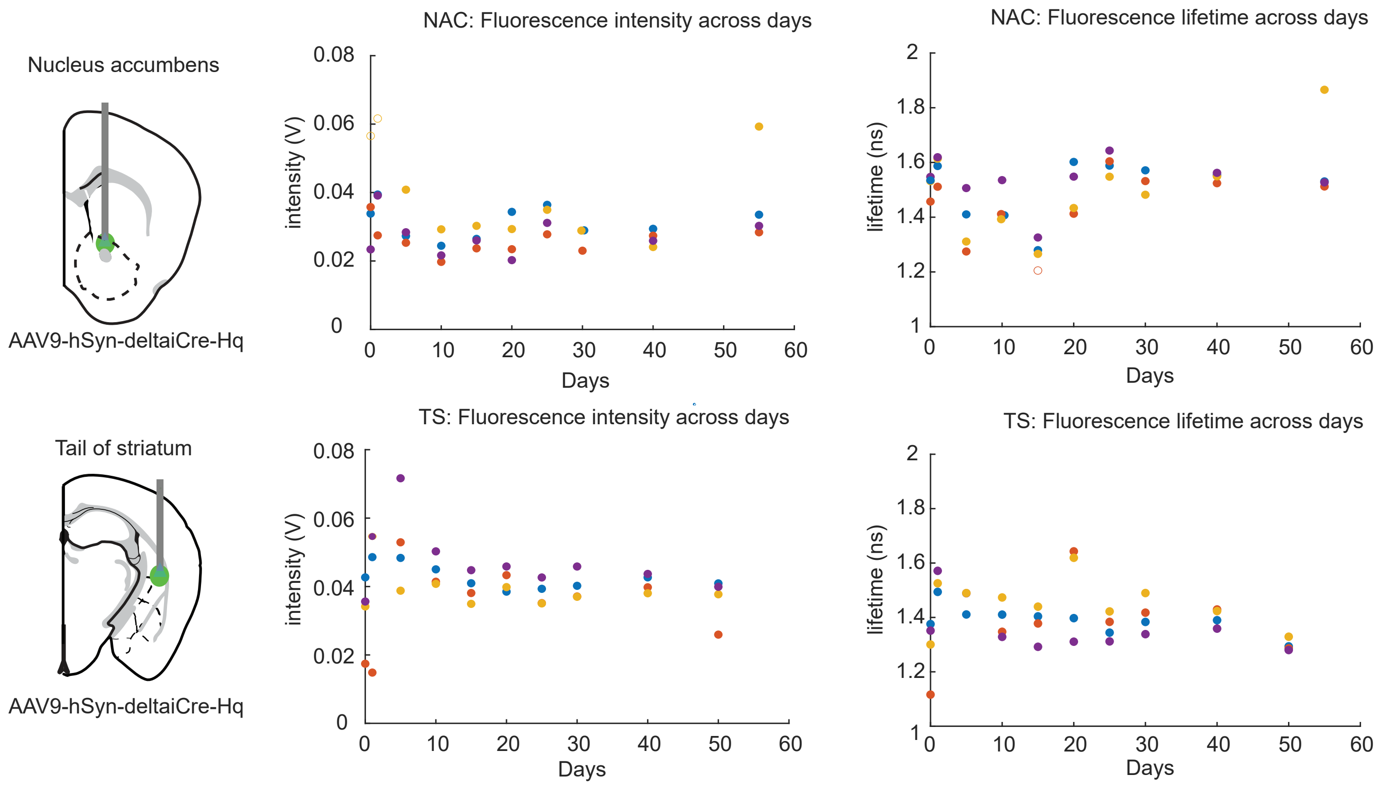
